# Allicin From Garlic Disrupts the Cytoskeleton in Plants

**DOI:** 10.1101/2021.03.08.434082

**Authors:** Ulrike Noll, Miriam Schreiber, Monika Hermanns, Christopher A. Mertes, Alan J. Slusarenko, Martin C.H. Gruhlke

**Affiliations:** Department of Plant Physiology, RWTH Aachen University, Worringer Weg 1, 52056 Aachen, Germany; College of Life Sciences, University of Dundee, Division of Plant Science, Errol Road, Invergowrie, Dundee, UK

**Keywords:** Allicin, Redox, *Tradescantia*, Arabidopsis, Cytoskeleton, Auxin

## Abstract

Allicin is a defence substance produced by garlic cells upon injury. It is a thiosulfinate showing redox-activity and a broad range of antimicrobial and biocidal activity. It is known that allicin efficiently oxidizes thiol-groups and it has been described as a redox toxin. In order to learn more about the effect of allicin on plants we used pure synthetized allicin, and investigated cytoplasmic streaming in sterile filaments of *Tradescantia fluminensis*, organelle movement using transgenic *Arabidopsis* with organelle-specifics GFP-tags, and effects on actin and tubulin in the cytoskeleton using GFP-tagged lines. Auxin distribution in roots was investigated using PIN1:GFP, PIN3:GFP, DR5:GFP and DII-VENUS Arabidopsis reporter lines.

Allicin inhibited cytoplasmic streaming in *T. fluminensis* and organelle movement of peroxisomes and the Golgi apparatus in a concentration-dependent manner, inhibited root growth and destroyed the correct root tip distribution of auxin.

We speculate that the cytoskeleton can be a primary “receptor” for allicin’s oxidizing properties and as a consequence cytoskeleton-dependent cellular processes are disrupted.

## Introduction

Actin and tubulin are major structural components of the cytoskeleton and their polymerization and depolymerization are important, amongst other things, for cytoplasmic streaming and intracellular trafficking (Allen & Allen, 1978). Soon after the discovery of actin it was shown that oxidizing agents not only inhibited the polymerization of globular actin, but also depolymerized actin filaments (Feuer et al. 1948). The importance of sulfhydryl groups for protoplasmic streaming in plant cells was reported in 1964 by Abe (Abe 1964) who showed the effect of the specific thiol-trapping reagent *p*-chloromercuribenzoate in bringing cytoplasmic streaming to a standstill in internodal cells of *Nitella flexilis*, leaf cells of *Elodea densa*, root hair cells of *Hydrocharis morsus ranae*, and stamen hair cells of *Tradescantia reflexa*. Elements of the cytoskeleton are known to be highly sensitive to oxidation (Farah and Amberg 2007; Farah et al. 2011) and redox-mediated posttranslational modifications of the cytoskeleton affect polymerization (Johansson and Lundberg 2007; Wang et al. 2001). The redox regulation of actin and tubulin dynamics is now a well characterized phenomenon (Wilson et al. 1916; Landino et al. 2004).

Allicin from garlic is the most prominent sulfur-containing bioactive compound in freshly damaged garlic tissue (Cavallito, 1944; Block 1992). It oxidizes thiol-groups in a thiol-disulfide-exchange-like manner leading to S-thioallylated adducts (Gruhlke et al. 2011; Gruhlke et al. 2010; Miron et al. 2010; Pinto et al. 2006; Winkler et al. 1992). Since allicin is a strongly antimicrobial compound with a broad-spectrum of activity, it has the potential for application in agricultural pest control (Auger et al. 2004; Freeman and Kodera 1995; Portz et al. 2008; Slusarenko et al. 2011; Slusarenko et al. 2008). Allicin’s degradation products, primarily polysulfanes, ajoenes or vinyldithiins, can also show biological activity and can be active as antimicrobial components. The effect of these sulfur-containing components in allelopathic interactions with other organisms can be of importance, as is observed in particular for wild garlic (*Allium ursinum* L.), attributable to the exudation of sulfur-containing secondary metabolites (Block 2010).

In animal cells it was shown that allicin, which acts as a “redox-toxin” (Gruhlke et al. 2010), affects the integrity and function of the cytoskeleton (Prager-Khoutorsky et al. 2007). It was reported that 0.5 μM allicin preferentially affected the tubulin cytoskeleton, while the actin cytoskeleton was a minor target (*ibid*.). However, at a higher 25 μM concentration of allicin the actin cytoskeleton was also reported to be disrupted (Sela et al. 2004). Furthermore, 16 cytoskeletal proteins were shown to be S-thioallylated by 100 μM allicin in human Jurkat cells within 10 minutes of exposure (Gruhlke et al. 2019).

The actin cytoskeleton is of particular importance for the correct polar transport of auxin in relation to PIN efflux carrier recycling (Vieten et al. 2007; Nick et al. 2009). Auxin also exerts an influence on the regulation of actin turnover (Rahman et al. 2007). The high redox sensitivity of actin is probably significant for the observation that auxin transport is related to the state of the total cellular glutathione pool, the most important thiol-based redox buffer of the cell (Koprivova et al. 2010). Thus, treatment of Arabidopsis seedlings with buthionine-sulphoximine (BSO), a specific inhibitor of glutathione biosynthesis, led to a clear concentration-dependent inhibition of root growth, and a loss of the auxin efflux carriers PIN1, PIN2 and PIN7 and, consequently, a destruction of the correct auxin distribution in the root tip (*ibid*.). This observation correlates with the finding that mutants in glutathione biosynthesis form shortened roots, such as is seen in the weak allele mutants *pad2* and *cad2* of the Arabidopsis *GSH1* gene which have reduced glutathione levels (Bashandy et al. 2010). Mutants of the strong *rml1* (root-meristemless) allele of *GSH1*, are almost unable to produce glutathione, are no longer able to maintain the root apical meristem, and consequently cannot form a root (Cheng et al. 1995; Vernoux et al. 2000).

Due to the need to understand the phytotoxicity of allicin with regard to a possible use in organic farming as well as the allelopathic effects of these substances, we are interested in understanding the molecular causes of the effects of allicin on plants. It has already been reported that allicin inhibits *Arabidopsis* primary and lateral root growth in a concentration-dependent manner (Borlinghaus et al. 2014, Leontiev et al. 2018). In the work reported here we analyzed the effect of allicin, applied as a solution of chemically synthesized pure allicin, on the cytoskeleton and, in consequence, cytoskeleton-dependent cellular processes like cytoplasmic streaming, organelle movement and auxin distribution which might lead to the root growth-inhibiting effects of allicin.

## Materials and Methods

### Plant material and cultivation

*Tradescantia fluminensis* V_ELL_. was cultivated in the greenhouse. For induction of flowering, the plants were grown in nutrient-poor soil.

Mutants and different GFP-tagged Arabidopsis lines used in this study are listed in Table 1. The plants were grown on solid Hoagland-medium (1 mM CaNO_3_; 5,1 mM KNO_3_; 0,5 mM MgSO_4_; 0,5 mM MgCl_2_; 0,13 mM (NH_4_)H_2_PO_4_; 30 μM NH_3_NO_3;_31 μM NaOH; 22 μM EDTA-Iron-Sodium Salt; 9,7 μM BH_3_O_3_; 22 μM FeSO_4_; 2 μM MnSO_4_; 0,31 μM ZnCl_2_, 0,21 μM CuSO_4_; 0,14 μM Na_2_MoO_4_; 86 nM Co(NO_3_)_2_, containing 8 g/L Plant Agar (Duchefa, Haarlem, The Netherlands). After stratification for two days at 4°C, the seedlings were grown under long day conditions (16 h dark, 8 h light; 22°C) for two days before treatment.

| mutant | Phenotype | Reference |
| --- | --- | --- |
| ABD2-GFP | GFP-fusion protein with actin binding protein ABD2 allows visualization of actin cytoskeleton | Voigt et al., 2005 |
| Tubulin-GFP | GFP-fusion with beta-tubulin allows visualization of tubulin cytoskeleton | Ueda et al., 1999 |
| GFP-PTS | Fusion of GFP with peroxisomal targeting sequence allows visualization of peroxisomes | Mano et al., 2002 |
| *pad2* | weak GSH1 allele, homozygote has ~30 % GSH content of the wild type | Glazebrook and Ausubel 1994 |
| *axr1-12* | No auxin-perception | Timpte et al., 1995 |
| *aux1-2* | Lack of auxin-importer AUX1-2. | Timpte et al., 1995 |
| *DR5::GFP* | GFP expression dependent on the auxin-responsive DR5-promoter. | Friml et al., 2003 |
| *DII-VENUS* | Degradation of the fluorescent protein depending on the presence of auxin | Brunoud et al., 2012 |

### Synthesis of allicin and quantification of allicin

Chemical synthesis of allicin as described previously (Gruhlke et al. 2010). Purity of chemically synthezised allicin was >95% (*ibid*.).

### Measurement of protoplasmic streaming

As a measure of the influence of allicin on protoplastic flow, the time was recorded after which cytoplasmic flow ceased on exposure to allicin. For this purpose, individual sterile flower filaments were plucked off with tweezers and microscoped as a water preparation. To prevent the specimen from being crushed, the cover glass was slightly raised by small amounts of plasticine at the corners. Treatment with allicin or garlic juice was carried out by using a filter paper to draw off the water under the coverslip and replacing it with allicin solutions of the appropriate concentration. In the same way, the allicin solution was exchanged for distilled water for the regeneration experiments. Microscopy was carried out at 400x magnification using transmitted light microscopy. Separate staining was not necessary.

### Confocal Laser Scan microscopy

For GFP-visualization, a Confocal Laser Scan Microscope (TCS SP1, Leica GmbH, Wetzlar, Germany) was used with an excitation wavelength of 488 nm and emission filter of 500 – 550 nm. For microscopy, a 63x water immersion objective was used and processed with ImageJ (http://rsbweb.nih.gov/ij).

For analysis of the cytoskeleton, two-day-old seedlings were used; the seedlings were placed on a slide in water or in an allicin solution of the indicated concentration. The analysis was carried out after 30 minutes. For the PIN and auxin localization study, seedlings were placed on Hoagland medium containing a concentration of 2 mM. The seedlings were incubated for 24 h under short day conditions (22°C, 16 h dark, 8 h light) before microscopic analysis.

### Growth assays

Arabidopsis seeds were surface sterilised with ethanol and spread with a toothpick on solid Hoagland medium. Plates were stored at 4°C for two days for stratification before incubation at 22°C for a further two days under long-day conditions (16 h light/8 h dark). After this time, seedlings were carefully transferred with forceps to Hoagland medium plates containing allicin at a final concentration of 0.5 mM and further incubated under these conditions. Root length was measured at day 0 and after three days.

### Statistics

Statistical analyses were carried out using the programme SigmaStat. Data were tested for statistical significance using a multivariate data analysis (ANOVA) with a significance level p<0.05.

## Results & Discussion

### Protoplasmic streaming in *Tradescantia fluminensis* V_ELL_

As a model system for the influence of allicin on the cytoskeleton, we chose the sterile hyaline filaments from the flower of *T. fluminensis* (shown in Figure 1 A, B & C). The time that elapsed until cytoplasmic streaming ceased after exposure to allicin solutions of various concentrationswas considered as a measure of the relative activity of allicin on the cytoskeleton. Particles carried in the cytoplasmic stream observed to follow locomotion (Fig. 1 D, E). Soon after the hyaline filaments of *T. fluminensis* were exposed the flow slowed down and stopped entirely after a certain time, which was dependent on the allicin concentration. As the concentration of allicin increased, the time required for the protoplasmic flow to come to a standstill decreased (Fig. 1D). While a concentration of 0.5 mM allicin led to a cessation of protoplasmic flow after approx. 13 minutes, a treatment with 4 mM allicin led to a complete cessation of protoplasmic flow after approx. 2 minutes (Fig. 1D).

**Figure 1:**
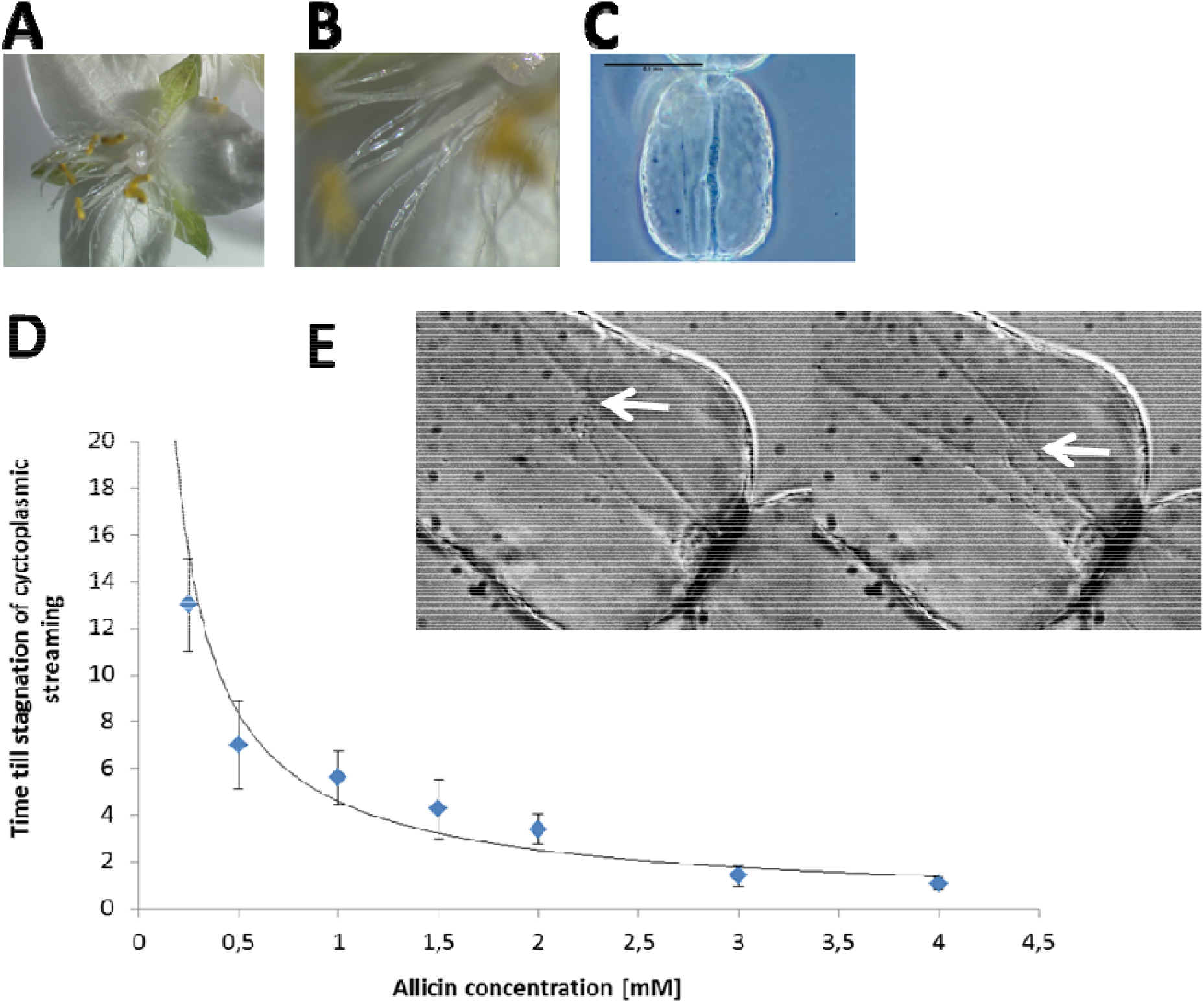
(A) (B) & (C) Sterile filaments in the flower of *Tradescantia fluminensis* are made up of hyaline cells and are well suited to observe protoplasmic streaming. The scale bar in (C) = 0.1 mm (D) Allicin leads to a concentration- and time-dependent decrease in protoplasmic flow. The time that elapses at a certain concentration of allicin until protoplasmic flow stops was measured, showing a correlation between the allicin concentration and the time that elapses until flow stops. (E) Movement of structures in the protoplasmic stream, in this case a fork in cytoplasmic strands, can be used to measure the protoplasmic streaming.

A further observation for 5, 10 and 30 minutes after treatment showed that in the presence of allicin no new onset of protoplasmic disturbance was observed.

### The effect of allicin on the protoplasmic streaming was partially reversible

In the following experiments it was investigated whether the effect of allicin was reversible by washing out the allicin. Filaments of *Tradescantia* were treated with allicin of a final concentration of 0.25 mM. After cytoplasmic flow stopped the allicin solution was washed by capillary suction with filter paper drawing in pure water at the coverslip edge to exchange the bathing solution. After about 12 minutes, protoplasmic flow began again in isolated cells, although at a slower rate than before allicin treatment (not shown). Thus, it can be concluded that the inhibition of protoplasmic flow by allicin can be reversible after a certain regeneration phase; at a concentration higher than 0.25 mM allicin, this reversal did not occur even after a long observation period (>30 minutes).

### Influence of allicin on the movement of peroxisomes in leaves

Since the movement of organelles, such as peroxisomes, also takes place within the framework of protoplasmic flow, an Arabidopsis line was used whose peroxisomes were tagged with GFP (Mano et al., 2002). This makes their movement traceable. Leaves of this Arabidopsis line were infiltrated by vacuum infiltration with either water as a control treatment or an allicin solution of the indicated concentration and the movement of the organelles was observed under the confocal laser scanning microscope (Fig. 2 A, B and C). While in the control it can clearly be seen by observing the curved trail of peroxisome movements between the arrows that the peroxisomes moved substantially over the 10 min observation period (Fig. 2 A), in the sample infiltrated with 4 mM allicin, movement of the peroxisomes was not seen (Fig. 2 B).

**Figure 2:**
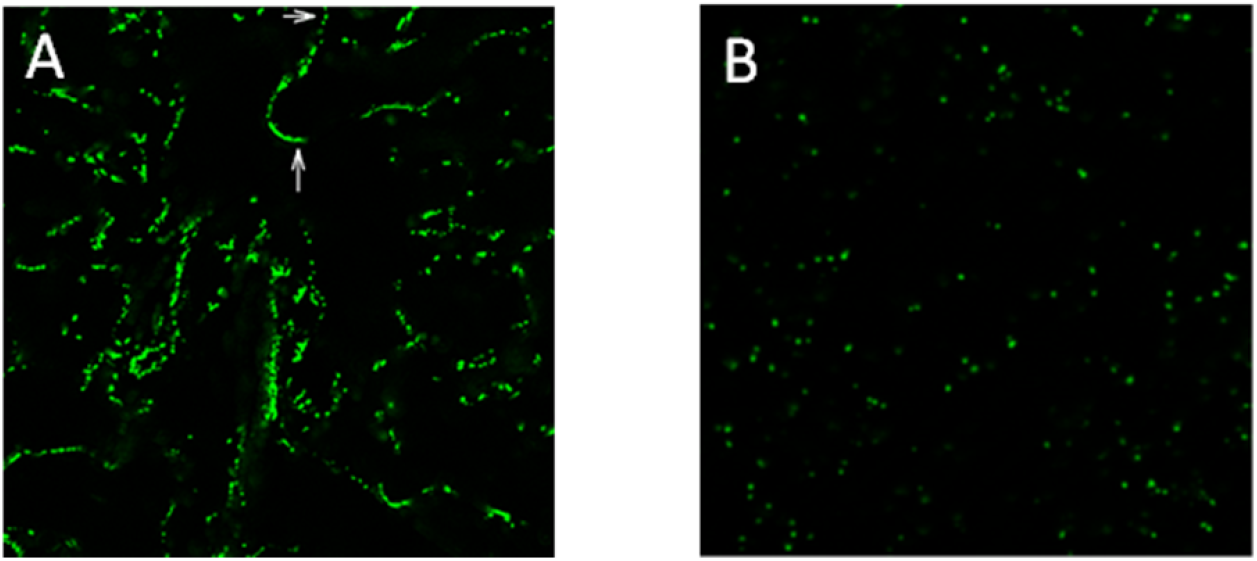
Movement of GFP-labelled peroxisomes in *Arabidopsis thaliana* treated with either (A) water as a control or (B) 4 mM allicin. The images show a stack of 20 images each, taken 30 seconds apart. (C) For reference, a single image of the control treatment is shown superimposed on the light microscopy image of the leaf cells.

### Allicin inhibits primary and lateral root growth and root hair development

The morphology of Arabidopsis wild type Col-0 seedlings germinated for two days then exposed to 0.5 mM allicin was very reminiscent of the morphology of the *rml1* mutant (Cheng et al. 1995). The roots of wild type Col-0 ceased growing in the presence of allicin, while the shoot axis appeared to be largely unaffected and developed normally (Fig. 3 A). Compared to the untreated control, the roots of plants exposed to allicin are generally much shorter and showed no lateral roots or root hairs. The prooxidant cumene hydroperoxide also led to a similarly shortened root (Fig. 3 B). Root growth in the *pad 2* mutant was also similarly inhibited by allicin and cumene hydroperoxide under the conditions tested (3B).

**Figure 3:**
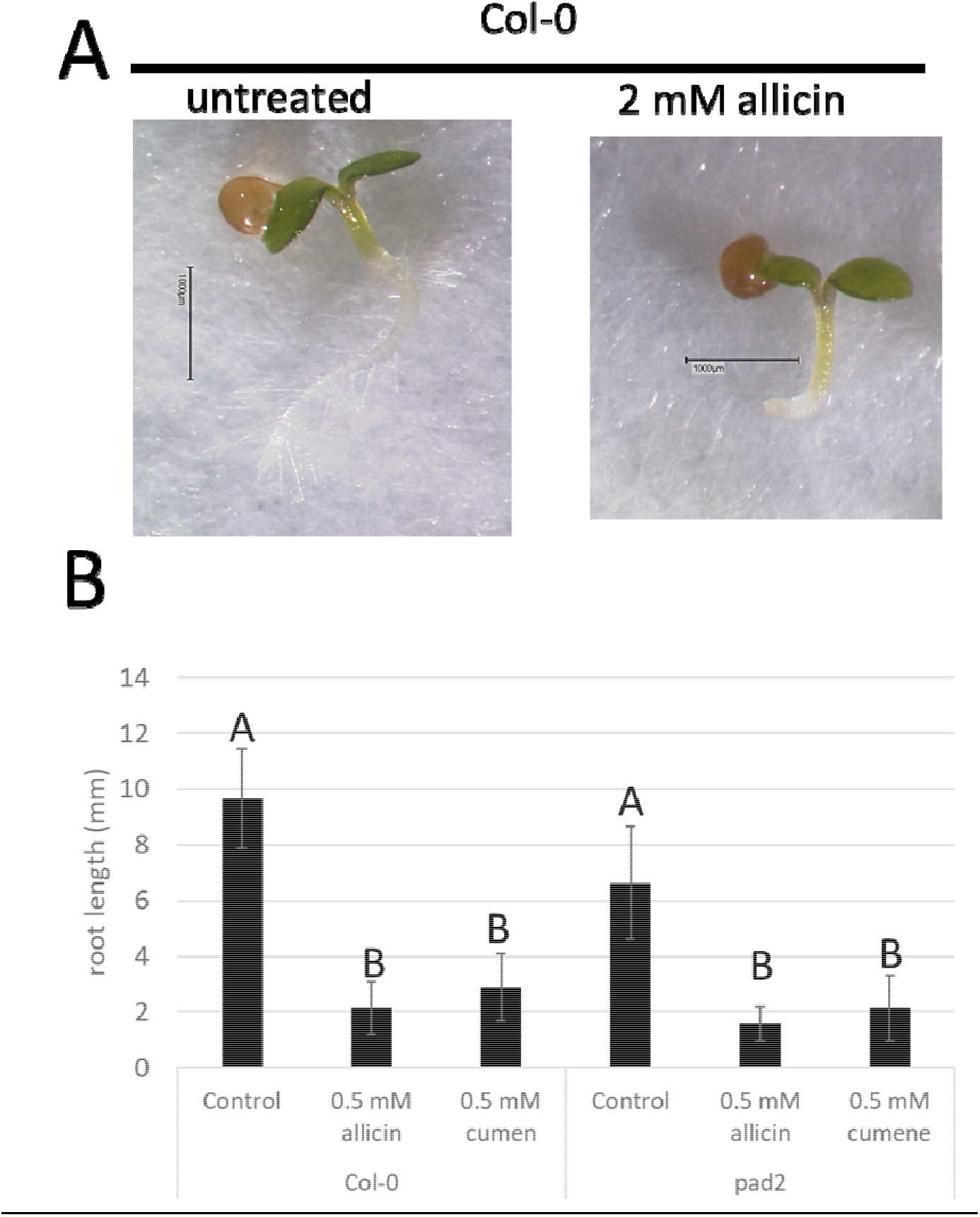
Allicin inhibits primary root growth and root hair development. Seeds were germinated for two days on filter paper discs which were then lifted and transferred to fresh medium containing allicin (A) or cumene hydroperoxide at the stated concentrations. (B) Wild type Col-0 was compared to the GSH-deficient *pad2* mutant. The scale bars = 1 mm. N=20, experiment was repeated three times with similar results.

### Effect of allicin on actin- and tubulin cytoskeleton in *Arabidopsis thaliana*

To understand in more detail how allicin affects root development and whether this is associated with the putative effect on the cytoskeleton based on the observation in *Tradescantia, Arabidopsis* lines were used whose actin or tubulin filaments have been visualised with by beta-tubulin:GFP fusion (Ueda et al., 1999) and GFP:actin-binding protein expression (Voigt et al., 2005).

Confocal microscopy of the actin cytoskeleton clearly shows typical filamentous structures in the root of the untreated control (Fig. 4 4 A). After exposure to 2 mM allicin for 24 h, the filamentous structures were destroyed; instead, GFP aggregate structures were visible at the margins of the cells. Allicin thus destroys the integrity of the actin cytoskeleton in roots of Arabidopsis.

**Figure 4:**
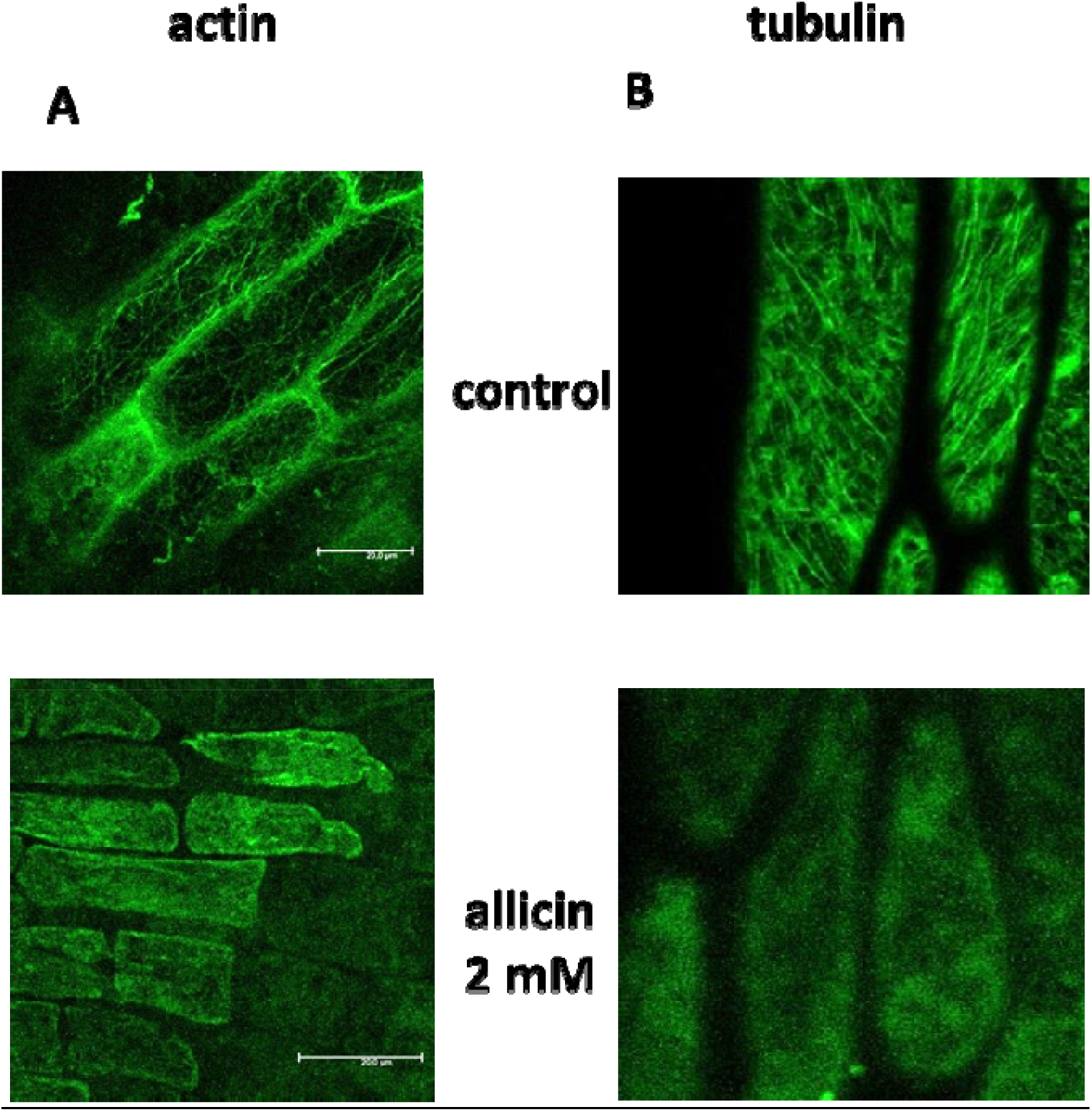
Effect of 2 mM allicin on the actin and tubulin cytoskeleton after exposure for 24 hours. While controls exposed to water only show a typical filamentous structure of both actin and tubulin, this is completely dissolved after 24 h exposure to allicin.

In addition to the actin cytoskeleton, the tubulin cytoskeleton was also examined in an analogous manner. Here, too, the typical filament pattern was seen in the untreated control; treatment with allicin resulted in a dissolution of these filament structures and a diffuse fine-particulate distribution of GFP fluorescence, in contrast to the aggregate formation observed for actin.

### Effect of allicin on auxin distribution PIN localization

The cytoskeleton fulfils a wide variety of tasks in the cell and is involved, among other things, in transport processes in the cell. In order to investigate the influence of cytoskeletal integrity - here primarily concerning actin - on important cellular transport processes, we chose the localisation of PIN proteins, whose localisation is crucial for regulated auxin transport in tissues. Again, GFP-tagged versions of the proteins, in this case PIN1 and PIN3, were used to track their localisation *in vivo*. In addition, the expression of the constructs is under the respective native promoter, so that the localisation of the expression can also be followed at tissue level.

In the untreated control, PIN1 shows expression mainly in the region of the central cylinder (Fig. 5 A); at the cellular level, the strongest fluorescence was observed on the basipetal side of the cell. Exposure to allicin of a concentration of 2 mM led to a complete loss of this clear localisation pattern and GFP fluorescence is weakly distributed throughout the cell (Fig. 5 B). This corresponds exactly to the observations made by Koprivova during treatment with BSO, suggesting that allicin is also active here via the glutathione pool (Koprivova et al., 2010).

**Figure 5:**
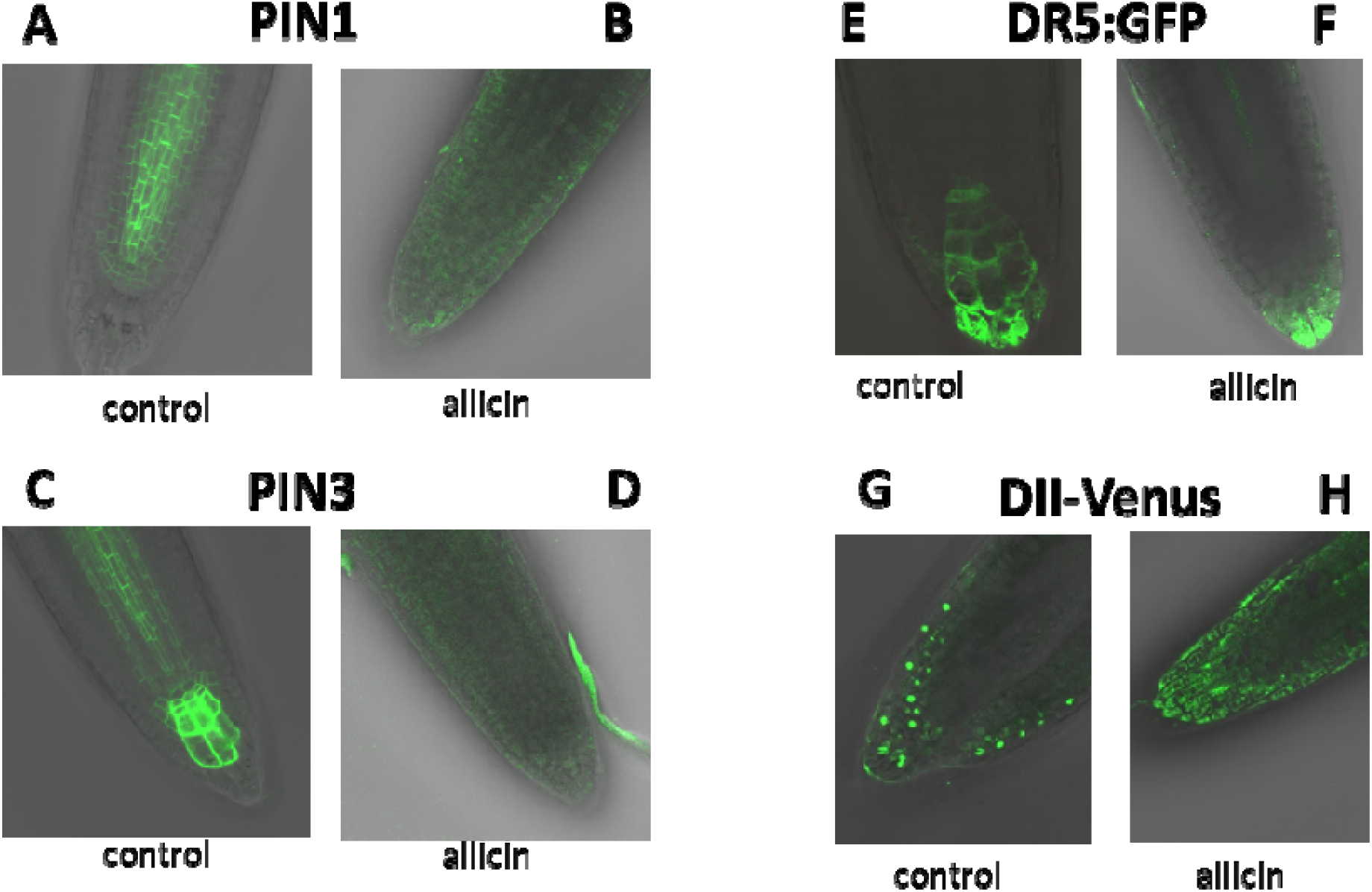
Influence of allicin on the localisation of PIN1 and PIN3 proteins and on auxin distribution in the root tip. (A) While the localisation of PIN1 in the control is restricted to the central cylinder, a diffuse localisation of PIN1 is seen in (B) allicin-exposed plants. (C) The localisation of PIN3 in the control is seen in the central cylinder and in the meristematic zone of the root tip, but under allicin exposure (D) the localisation of PIN 3 protein is also diffuse. This is also reflected in the localisation of auxin. In control plants, where GFP expression is under the auxin-responsive DR5 promoter, a typical distribution of auxin is seen in the root tip, meristematic and quiescent zone (E), whereas this clear localisation is abolished under allicin exposure (F). The detection of auxin distribution in the root by means of DII-venus indicates in a complementary way that no auxin is present where the fluorescent protein is observed, because auxin leads to the degradation of the protein. Accordingly, in the control (G), fluorescence is seen in the marginal areas of the root, but not in the meristematic and quiescent zone, whereas under allicin exposure, fluorescence is observed everywhere in the cells (H). All treated samples were exposed to 2 mM allicin for 24 h.

In contrast to PIN1, the localisation of PIN3 is also found in the area of the cortex, but it is more prominent in the area of the root apical meristem. Again, fluorescence is strongest at the basipetal end of the cell (Fig. 5 C).

As observed with PIN1, complete disruption of PIN localisation has also occurred with PIN3 by treatment with 2 mM allicin. The GFP signal, which indicates the localisation of the PIN1 protein, is diffusely present at both the cellular and tissue levels (Fig. 5 D). These results exactly mirror the effects of the GSH1 inhibitor buthione sulfoximine (BSO) on PIN distribution (Kopriva et al., 2010). BSO leads to a reduction in cellular GSH levels and this mirrors the result of allicin titrating out and oxidizing the GSH pool (Gruhlke et al., 2010).

### Effect of allicin on auxin distribution

Based on the observation that allicin destroys the localisation of PIN proteins, and because PIN proteins are essential for the directional transport of auxin, the final aim was to investigate whether aberrant localisation of auxin occurs after allicin treatment. Two independent methods based on fluorescent proteins were used to check the localisation of allicin. The DR5::GFP reporter system is based on the fact that auxin induces the activity of the DR5 promoter and thus drives GFP expression. GFP fluorescence thus indexes the presence of auxin.

Untreated Arabidopsis roots show a clear localisation of auxin around the quiescent zone of the root apical meristem, in the central cylinder and the epidermal layers of the root (Fig. 5E). Treatment with allicin resulted in the localisation of GFP fluorescence being restricted to the root apex; auxin-dependent expression of GFP is no longer found in the quiescent and division zone of the root apex (Fig. 5F). This shows that allicin causes a clear redistribution of auxin localisation at the root tip.

As an alternative method to detect the localisation of auxin at the tissue level, the DII-Venus construct was used, in which a “fast maturing” form of yellow-fluorescent protein (YFP) is expressed in frame with an IAA protein under the control of a constitutive promoter. Because IAA proteins are degraded in the presence of auxin, a lack of fluorescence of the VENUS protein indicates a high concentration of auxin, whereas a strong fluorescence of the protein indicates a low concentration of auxin. The results obtained by means of DII Venus corresponded to the statements we also perceived with DR5:GFP. While the control showed that auxin was present in the root meristem area, the distribution was diffuse after allicin exposure (Fig. 5G).

### Effect of allicin on mutants impaired in auxin homeostasis and signaling

Because we observed an effect on auxin distribution and PIN localization after allicin treatment (Fig. 5), we investigated the effects of allicin on the auxin transport mutant *aux1-2* and the auxin resistant mutant *axr1-12* in comparison to wt Col-0. Whereas the PIN proteins are functional in auxin efflux at the basal cell pole, the AUX1 protein transports auxin into the cell from the apoplast at the apical pole.

The Col-0 wild type shows that its exposure to 0.5 mM allicin leads to a significantly shortened root. The aux1-2 mutant, already has a significantly shortened root in the control compared to the Col-0 but in contrast, root length was not further affected by the presence of allicin, *i.e*. this mutant showed a degree of allicin resistance (Fig. 6 A) and this suggests that the root phenotype we observe with allicin may be dependent upon auxin perhaps over-accumulating to supraoptimal levels and inhibiting root growth. The *axr1-12* mutant, on the other hand, shows no significant change in root length phenotype compared to the wild type in the control, but treatment with allicin causes the roots to be shortened, but to a much lesser extent than in the Col-0 control. This suggests that the influence of allicin on auxin transport is partly responsible for the observation that roots grow less well under allicin exposure (Fig. 6 B). Possibly, another auxin receptor can partially complement the phenotype of *axr1-12*. To test this hypothesis, Col-0 and *axr1-12* mutants were exposed to 20 μM cytochalasin, which leads to disruption of the actin cytoskeleton. As a control, it was examined whether the solvent DMSO at the concentration used (1% v/v) would lead to an effect, which was not the case. While the Col-0 wild type showed significantly shortened roots after treatment with cytochalasin, the *axr1-12* mutant does not, suggesting that it is disruption of the actin cytoskeleton alters auxin localisation. Accordingly, a mutant that is not an auxin receptor mutant shows no phenotype in terms of root length upon destruction of the actin cytoskeleton. These results suggest that the effect allicin has on root development correlates, at least in part, with its influence on auxin distribution in the root dependent upon allicin’s effects on the cytokeleton.

**Figure 6:**
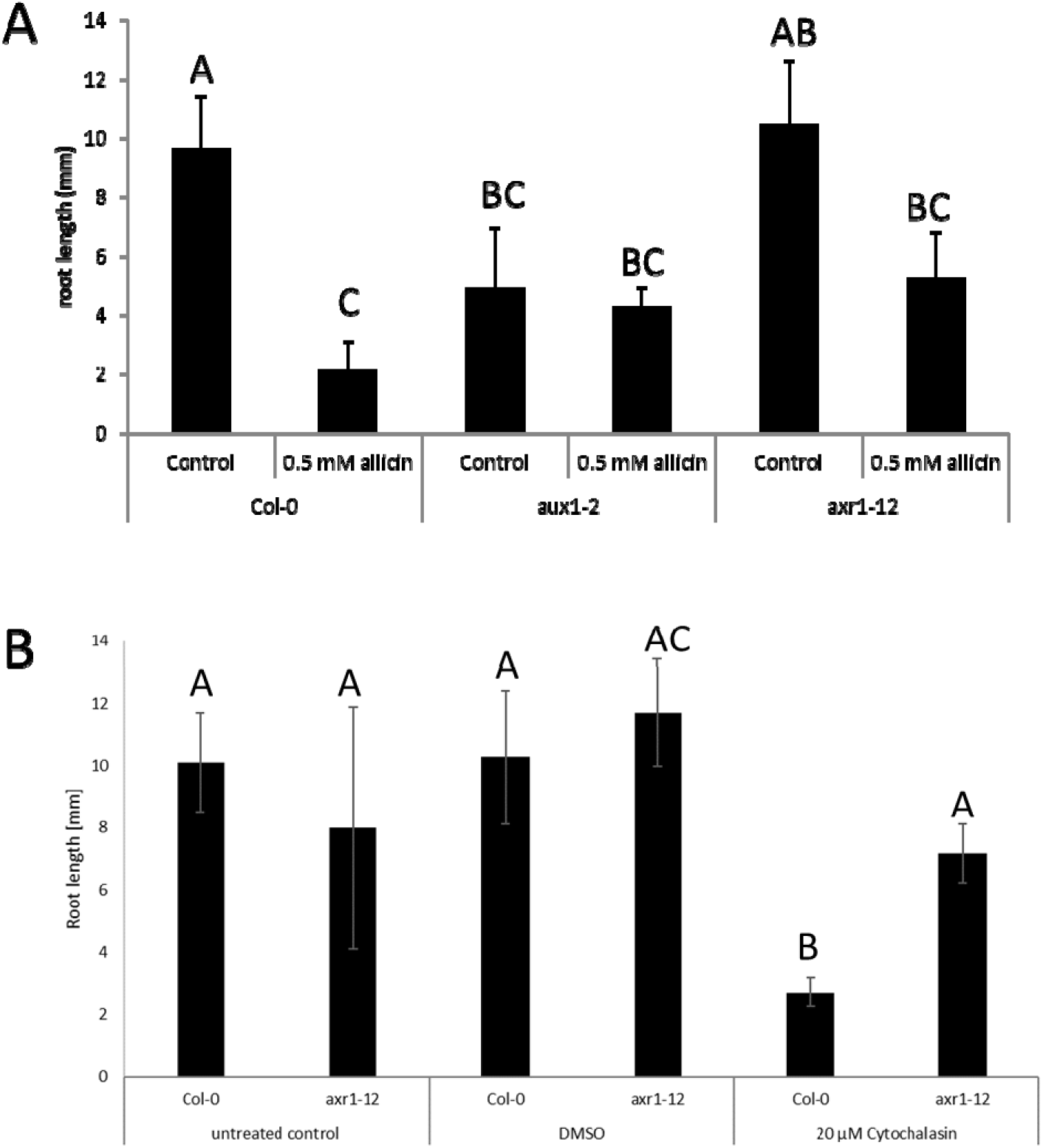
(A) Influence of allicin on *aux1-2* and *axr1-12* mutants. (n=20) the experiments were repeated three times with similar results.

## Supporting information

Control

Treated with allicin

## Acknowledgement

Financial support from RWTH Aachen University is acknowledged.

## Supplemental Material

Videos showing the inhibitory effect of allicin on the movement of GFP-tagged peroxisomes in Arabidopsis leafs.

## Notes

### Competing Interest Statement

The authors have declared no competing interest.

